# Differential responses of antioxidants and dehydrin expression in two switchgrass (*Panicum virgatum*) cultivars contrasting in drought tolerance

**DOI:** 10.1101/486308

**Authors:** Yiming Liu, Guofu Hu, Guoqiang Wu, Guodao Liu, Hengfu Huan, Xipeng Ding, Linling Yan, Xinyong Li, Bingyu Zhao, Xunzhong Zhang

## Abstract

Drought stress is a major limiting factor for plant growth and development in many regions of the world. This study was designed to investigate antioxidant metabolism and dehydrin expression responses to drought stress in two switchgrass cultivars (drought tolerant Alamo, and drought sensitive Dacotah) contrasting in drought tolerance. The plants were subjected to well-watered [100% evapotranspiration (ET)] or drought stress (30%-50% ET) conditions for up to 24 d in growth chambers. Drought stress decreased leaf relative water content (RWC), increased leaf electrolyte leakage (EL), leaf malondialdehyde (MDA) content in two cultivars, but Alamo exhibited higher leaf RWC level, lower leaf EL and MDA when compared to Dacotah at 24 d of drought treatment. Drought stress also increased superoxide dismutase (SOD), catalase (CAT) and ascorbate peroxidase (APX) activities in two cultivars, Alamo had relatively higher SOD, CAT and APX activities and greater abundance of SOD and APX isozymes than Dacotah at 24 d of drought treatment. Alamo had higher abundance of 55 KDa and 18 KDa dehydrin accumulation than Dacotah under drought treatment. Relative genes expression level of *PvCAT1, PvAPX2, PvERD and PvPIP1;5* in Alamo were significantly higher than Dacotah at 24 d of drought treatment. These results suggest that increase in antioxidant enzymes and accumulation of dehydrin were highly related with switchgrass drought tolerance. Antioxidant enzyme activity, isozyme expression and dehydrin abundance could provide a useful screening tool to identify relative drought tolerance in switchgrass cultivars.

## Introduction

Drought occurs in all climates and many parts of the world every year when sufficient water needed to sustain an area is not available, causing significantly impacts on plants growth, development and crop yield [1]. Worldwide losses in crop yields from drought probably exceed the losses from all other abiotic stress combined [2, 3]. Drought stresses cause oxidative stress by contributing to reactive oxygen species (ROS) such as superoxide radical (O^•–^_2_), hydrogen peroxide (H_2_O_2_), hydroxyl radical (OH•) and singlet oxygen (^1^O_2_) [4]. ROS can seriously disrupt normal metabolism through oxidative damage to nucleic acids, lipids, protein and damage membrane function [5]. However, ROS may also serve as a transduction signal during drought stress and improve stress defense mechanisms of plant [6]. ROS levels that are too low or too high affect plant growth and development, maintaining ROS levels within a moderate range is important for plants [7, 8].

Plants have developed an antioxidant defense system in response to the high level of ROS [9]. There are generally two repairing mechanisms that plants have developed to scavenge free ROS: (i) production of antioxidants or antioxidant enzymes that directly react with and scavenge ROS, including superoxide dismutase (SOD), catalase (CAT), ascorbate peroxidase (APX), peroxidase (POD), and a tocopherol; (ii) production of enzymes that regenerate oxidized antioxidants such as glutathione, glutathione reductase, ascorbate, and ascorbate reductase[10]. Previous studies provided correlative evidence that the enhanced drought tolerance of plant was associated with changes in antioxidant enzymes (SOD, POD, CAT, POX and GR) and maintenance of low H_2_O_2_ levels [11–13]. It is known that organelles with a highly oxidizing metabolic activity or an intense rate of electron flow, such as chloroplasts, mitochondria, and microbodies, are major sources of ROS, different isoenzymes such as Cu/Zn-SOD, Fe-SOD, Mn-SOD, cytosol APX (cAPX), and microbody APX (mAPX) have been found in different organelles [14]. The antioxidant isozymes could be used as a biochemical marker to study the tolerance of plant to stress. Sen and Alikamanoglu [15] found that two new POX isozyme bands were detected in all drought-tolerant sugar beet mutants compared to the control, and the intensity of Fe-SOD, Cu/Zn-SOD, CAT and APX isozymes were detected at different intensities among the drought-tolerant sugar beet mutants. Recent study found that a new CuZn SOD iszyme OsCSD3 which encoded by LOC_Os03g11960 of rice, was up-regulated in response to drought, oxidative stress and salt [16].

Dehydrin (DHN) is a multi-family of proteins present in plants that are produced in response to cold and drought stress. DHNs are hydrophilic and reliably thermostable. They are stress proteins with a high number of charged amino acids that belong to the Group II Late Embryogenesis Abundant (LEA) family [17, 18]. DHNs have been divided into five subclasses based on their conserved amino acid sequences: the Y, S and K segments and include YnSKn, SKn, Kn, YnKn and KnS sub-types[19]. It has been reported that the expression of DHN is positively correlated with the tolerance to cold, drought, and salt stress [20]. DHNs play an important protective role during cellular dehydration. The accumulation of DHNs was observed in roots, leaves, coleoptiles, crowns and seeds under drought stress [21].

Switchgrass (*Panicum virgatum*) was selected as a model bioenergy crop in the United Sates [22]. To avoid competition of arable lands with food crops, switchgrass will be mainly grown on marginal lands, where millions of hectares of these lands are drought-affected [23]. Two distinct switchgrass ecotypes are generally defined based on morphological characteristics and habitat: lowland and upland. Lowland ecotypes are mostly tetraploid (2n = 4× = 36) while upland ecotypes are mainly octaploid (2n = 8× = 72) or hexaploid (2n = 6× = 54) [24]. Lowland ecotypes are taller and higher biomass yield, whereas upland ecotypes are shorter and less biomass yield [25]. Improving switchgrass yield under drought stress is one of the most important goals of plant breeding. Our previous study have found that switchgrass exhibits a wide range of genetic variability in drought tolerance, Alamo ranked # 4 in drought tolerance and Dacotah ranked # 48 among 49 switchgrass lines [26]. However, limited information is available on the gene expressions in conjunction with antioxidant and dehydrin responses and isozyme alterations under drought tolerance in switchgrass. Identifying and understanding the function of antioxidant defense mechanisms are important for developing drought tolerant switchgrass plants. The objectives of this study were to determine whether drought resistance of switchgrass cultivars could be associated with antioxidant metabolism and dehydrin expression levels.

## Materials and methods

### Plant materials and culture

Two cultivars Alamo and Dacotah were examined in this study. Each switchgrass line was propagated by splitting tillers on Mar, 10, 2014. Five tillers from each line were transplanted into plastic pots (17 cm diam., 20 cm high, with four holes at the bottom for drainage) filled with 3.5 kg of a soil and sand mixture (soil:sand = 2:1 v/v, sand: 0.1-1.0 mm diam.). The plants were grown in greenhouse with temperatures of 30±1 °C/25±1 °C (day/night), a 14-h photoperiod, 75% relative humidity, and with photosynthetically active radiation (PAR) of approximately 500 μmol m^-2^s^-1^ (natural daylight supplemented with fluorescent lamps). The plants were irrigated daily, and fertilizer containing N (Bulldog brand, 28-8-18, 1% ammonia N, 4.8% nitrate N, and 22.2% urea N; SQM North America, Atlanta, GA) and micronutrients was applied at 0.1 lb. 1000 ft^-2^ every week.

After the plants were grown for two months and had reached the E5 developmental stage [27], then moved into a growth chamber for the experiment. The chamber was set at 30/25 °C (day/night temperature), 75% of relative humidity, 14 h photoperiod, and PAR at 500 μmol m^-2^s^-1^. Plants were fertilized once a week, and watered every two days until water drainage occurred at the bottom of the pot at each irrigation.

### Drought stress treatment

In order to determine the soil water content (SWC) of each pot more quickly, an equation of linear regression between the SWC and volumetric soil moisture content (VWC) was made before the drought treatment. Soil of eight pots was oven dried at 105°C for 48 h to obtain their dry weights (DW). Then we added enough water to each pots, after 1 h when no water leaked from the bottom of the pots, fresh weight of each pot and VWC were measured with a soil moisture meter (model HH2, Delta-T Devices, Cambridge, England) and every three day thereafter. SWC was determined using the formula: SWC (%) = (FW−DW)/DW×100. Then we got an equation of linear regression between the SWC and VWC (Supplemental table 1).

Plants were allowed to acclimate to growth chamber conditions for one week before drought treatments were imposed. Each line were randomly assigned to either the control group (n=4), which was kept well-watered (100% ET), or to the drought treatment group (n=4), in which the soil moisture was allowed to progressively decline from 0 d to 24 d. Each pots of drought treatment were weighted every two days and VWC were also collected, then SWC was calculated by the equation we got above (Supplemental table 1), the water needed to add of each pot was calculated to compensate for 30%-50% ET during the experiment over the 24 d period (Fig. 1).

**Fig. 1.**
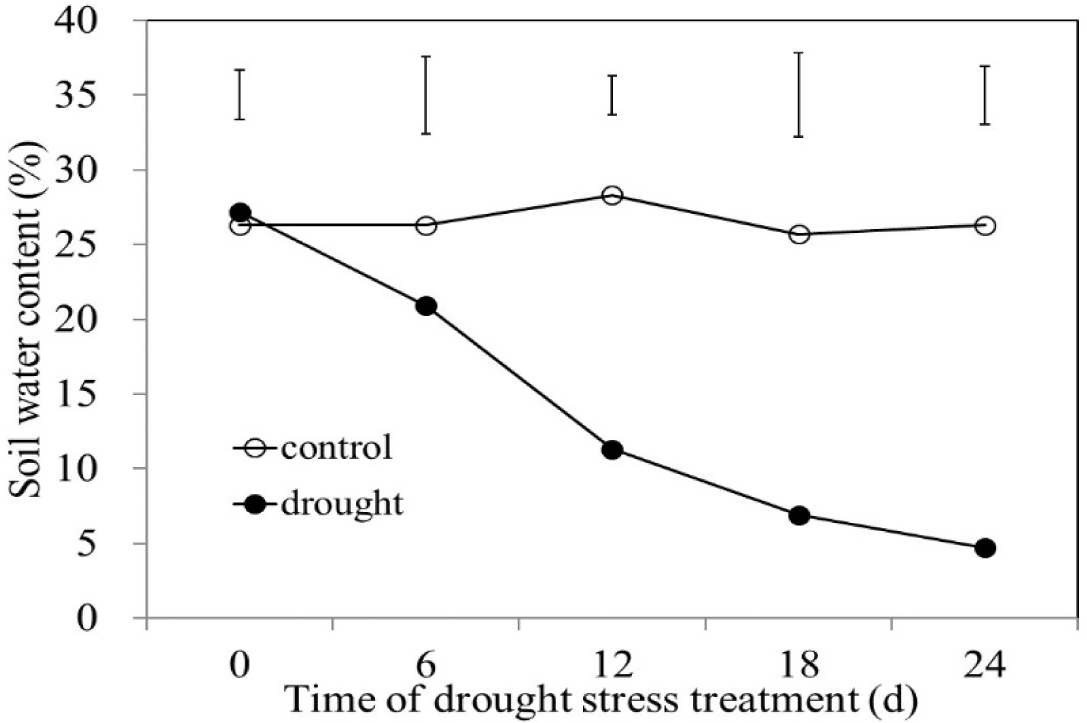
Effects of drought stress on soil water content (SWC). Vertical bars indicate LSD value (P=0.05).

### Physiological measurements

Leaf samples were collected for electrolyte leakage (EL) and relative water content (RWC) measurements at day 0, 6, 12, 18 and 24 of drought stress. Leaf tissues were also sampled, frozen with liquid N, and used for antioxidant emzyme and dehydrin analysis.

Leaf electrolyte leakage (EL) was measured according to the method of Marcum [28] with some modifications. The top 2^nd^ or 3^rd^ mature leaf blades were excised and cut into 2 cm segments. After being rinsed 3 times with deionized H_2_O, 0.2 g leaf segments were placed in a test tube containing 20 mL deionized H_2_O. The test tubes were agitated on a shaker for about 24 h and the solution conductivity (C_1_) was measured with a conductivity meter (SR60IC, VWR, Radnor, PA). Leaf samples were then autoclaved at 120 °C for 30 min, and the conductivity of the solution containing killed tissue was measured once the tubes were cooled down to room temperature (C_2_). The relative EL was calculated using the formula: EL (%) = (C_1_/ C_2_) ×100.

Leaf relative water content (RWC) was determined according to the method of Barrs and Weatherley [29]. The Leaf RWC was calculated based on the following formula: RWC= (FW-DW)/ (TW-DW) ×100, where FW is leaf fresh weight, DW is the dry weight of leaves after drying at 85 °C for 3 d, and TW is the turgid weight of leaves after soaking in distilled water for 24 h at 20 °C.

### Antioxidant enzyme activity

Frozen leaf samples were ground in liquid nitrogen and homogenized in 50 mM sodium phosphate buffer (pH 7.0) containing 2 mM EDTA, 5 mM β-mercaptoethanol and 4% (w/v) polyvinylpyrrolidone-40 (PVP-40). The homogenate was centrifuged at 12,000×g for 30 min at 4 °C. The supernatant was used for assay of the antioxidant enzymes CAT, APX, and SOD.

Total SOD activity was determined by measuring its ability to inhibit the photochemical reduction of nitro blue tetrazolium (NBT) according to the method of Giannopolitis and Ries [30] with minor modifications. The reaction solution (1 mL) contained 50 mM phosphate buffer (pH 7.8), 0.1 mM EDTA, 13 mM methionine, 65 μM NBT and 1.3 μM riboflavin, and 30 μL SOD extract. A solution containing no enzyme solution was used as the control. Test tubes were irradiated under fluorescent lights 60 μmol·m^-2^·s^-1^) at 25 °C for 10 min. The absorbance of each solution was measured at 560 nm using a spectrophotometer, and one unit of enzyme activity was defined as the amount of enzyme that would inhibit 50% of NBT photoreduction.

The CAT activity was determined using the method of Chance and Maehly[31] with modifications. For CAT, the reaction solution (1 mL) contained 50 mM phosphate buffer (pH 7.0), 15 mM H_2_O_2_, and 30 μL of extract. The reaction was initiated by adding the enzyme extract. Changes in absorbance at 240 nm were read in 1 min using a spectrophotometer (ϵ = 39.4 M^-1^ cm^-1^).

The APX activity was assayed by recording the decrease in absorbance at 290 nm for 1 min. The 1.5-mL reaction mixture contained 50 mM potassium phosphate buffer (pH 7.0), 0.5 mM ascorbic acid, 0.1 mM EDTA, 0.1 mM H_2_O_2_, and 0.15 mL of enzyme. The reaction was started with the addition of 0.1 mM H_2_O_2_ [32].

### Lipid peroxidation

Lipid peroxidation was measured in term of leaf MDA content [33]. A 1 ml aliquot of supernatant was mixed with 4 mL of 20% trichloroacetic acid containing 0.5% thiobarbituric acid. The mixture was heated at 100 °C for 30 min, quickly cooled, and then centrifuged at 10000 *g* for 10 min. The absorbance of the supernatant was read at 532 nm. The unspecific turbidity was corrected by A600 subtracting from A530. The concentration of MDA was calculated using an extinction coefficient of 155 mM^−1^ cm^−1^.

### Antioxidant isozymes

The procedure of protein extraction was the same as for antioxidant enzymes. The extracts (15 μL) for SOD and APX were loaded on each gel. Native polyacrylamide gel electrophoresis (PAGE) was performed using a Mini-Protean system (Bio-Rad Laboratories, Hercules, CA) at 4 ᴼC, 120 V for 90 min, except that SDS was omitted. For SOD and APX, enzyme extracts were subjected to native PAGE with 10% resolving gel and 3% stacking gel and CAT was detected on 7% resolving gel and 4% stacking gel.

The total activity of SOD was revealed using the method of Beauchamp and Fridovich[34] with some modifications. The gels were incubated in 50 mM potassium phosphate buffer (pH 7.5) containing 2.5 mM NBT in dark for 25 min. After being washed twice with the same buffer, the gels were soaked in 50 mM potassium phosphate buffer (pH 7.5) containing 30 μM riboflavin and 0.4% N,N,N,N-tetramethylethylenediamine 235 (TEMED) in the dark for 40 min. The gels were then illuminated for 10 min with gentle agitation until appearance of enzyme bands and were transferred to 1% (v/v) acetic acid to stop the reaction.

The activity of APX was detected using the method of Lopez-Huertas *et al*. [35] with some modifications. The gels were pre-incubated in 50 mM sodium phosphate buffer with 4 mM ascorbate and 2 mM H_2_O_2_ for 20 min. After briefly being washed with 50 mM potassium phosphate buffer (pH 7.0), the gels were stained in 50 mM potassium phosphate buffer (pH 7.8) containing 28 mM TEMED and 1.25 mM NBT until the bands were clearly visible. The gels were then washed with distilled water to stop the reaction.

### SDS-PAGE and western blot

Frozen leaf samples were ground in liquid nitrogen and re-suspended in100 μl 3 × Laemmli buffer containing 16% β-mercaptoethanol.The tissue was then boiled for 10 min and pelleted at a high speed for 10 min.Twenty micro liters of protein extract was applied to and separated on a 10% SDS-PAGE gel. The proteins were blotted to a PVDF membrane using a Bio-Rad Trans-Blot R TurboTM Transfer System. The membrane was blocked with 5% nonfat skim milk in 1 × Tris-saline buffer supplemented with 0.5% Tween20 (1 × TBST). After a brief rinse with TBS, the membrane was incubated in TBS with a dehydrin polyclonal antibody raised from rabbit (Assay Designs) at a dilution of 1:250 for 1.5 h. Next, the membrane was rinsed in TBS containing 0.5% Tween 20 (TBS-T) four times and then placed for 1 h in a solution of goat antirabbit IgG (dilution 1:17500) conjugated to alkaline phosphatase (Sigma). The membrane was rinsed in TBS-T four times.The chemiluminescent signals were exposed to autoradiography film (Genesee Scientific, SanDiego, CA) using a Kodak film processor SuperSignal West Pico Chemiluminescent Sbustrate (Prod # 1856136, Thermo Scientific).

### RNA extraction and quantitative reverse transcription PCR (qRT-PCR)

Total RNA was extracted from 150 mg of leaf tissue using RNeasy plant mini RNA kit (50) (Qiagen, Valencia, CA), and RNA samples were further treated with DNase (Promega, Madison, WI, USA) to eliminate DNA contamination. Integrity of RNA was confirmed with miniaturized gel electrophoresis with the Agilent Bioanalyzer (Agilent Technologies, Inc., Santa Clara, CA). cDNA was synthesized using the DyNAmo cDNA Synthesis Kit (New England Biolabs, Ipswich,MA, USA). The qRT-PCR analysis primer pairs of the corresponding genes were designed according to sequences obtained from the Phytozyme website (Supplemental table 2). Each 20-μl reaction, which contained 15 ng of random hexamers/μl, 10 IU of Moloney murine leukemia virus RNase H^+^ reverse transcriptase solution (Thermo Fisher Scientific), and appropriate buffer containing deoxynucleoside triphosphates (dNTPs) and MgCl_2_ in a final concentration of 5 mM (1×; Thermo Fisher Scientific), was incubated at 25°C for 10 min and at 37°C for 30 min, inactivated at 85°C for 5 min, and finally chilled to 4°C. Two replicate RT reactions were made for each RNA sample.

### Experimental design and statistical analysis

The experiment was a 4×2 factorial combination (two switchgrass cultivars, and two drought levels: well-watered and drought treatment) in a complete block design (one treatment of one species served as the block) with four replications. All data were subjected to analysis of variance (ANOVA, SAS 8.1, SAS Institute Inc., Cary, NC). The treatment means were separated using Fisher’s protected least significant difference (LSD) test at 5% probability level.

## Results

### Effects of drought stress on physiological parameters

Drought stress reduced leaf RWC, increased leaf EL and MDA regardless of cultivars (Fig. 2, 3 & 4). There were significant differences in leaf RWC, leaf EL and MDA in the two cultivars under drought stress conditions when compared to the controls at 18 d and 24 d. Leaf RWC, leaf EL and MDA of Dacotah decreased or increased sharply at both 18 d and 24 d of drought treatments. Alamo had a relatively higher leaf RWC and lower leaf EL and MDA than Dacotah at 12 d, 18 d and 24 d of drought stress.

**Fig. 2.**
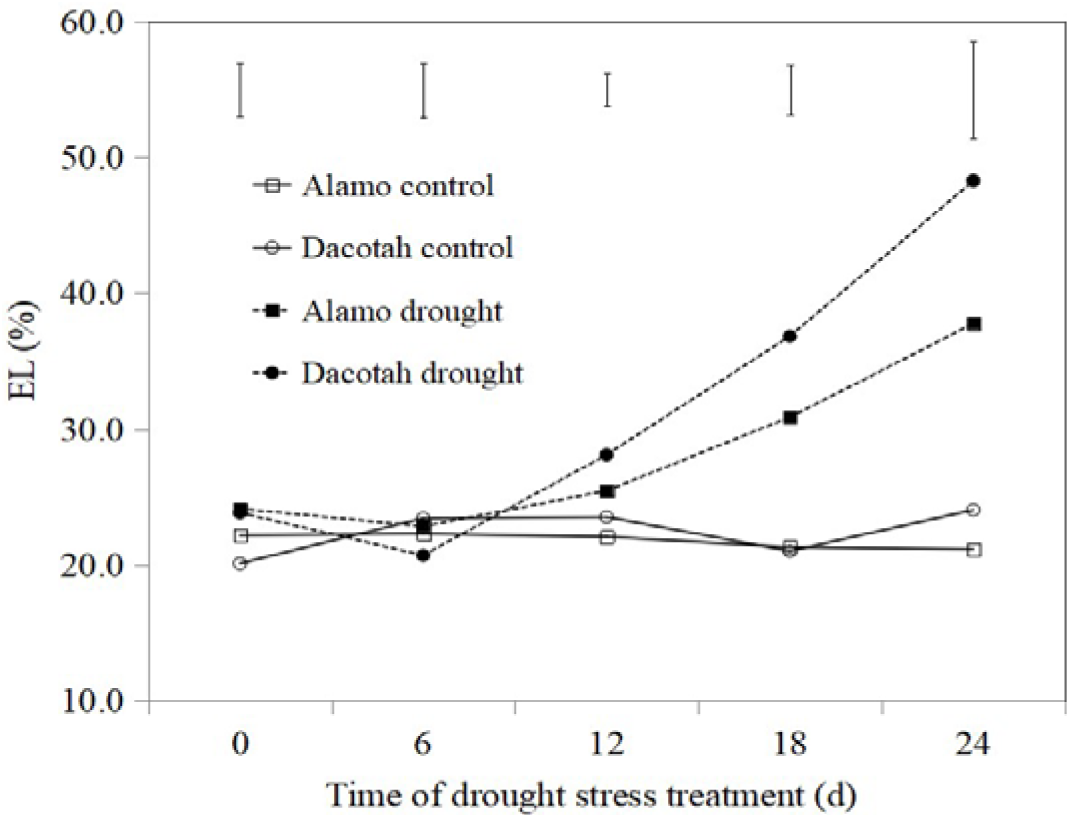
Effects of drought stress on leaf electrolyte leakage (EL) of two switchgrass cultivars Alamo and Dacotah. Vertical bars indicate LSD value (P=0.05).

**Fig. 3.**
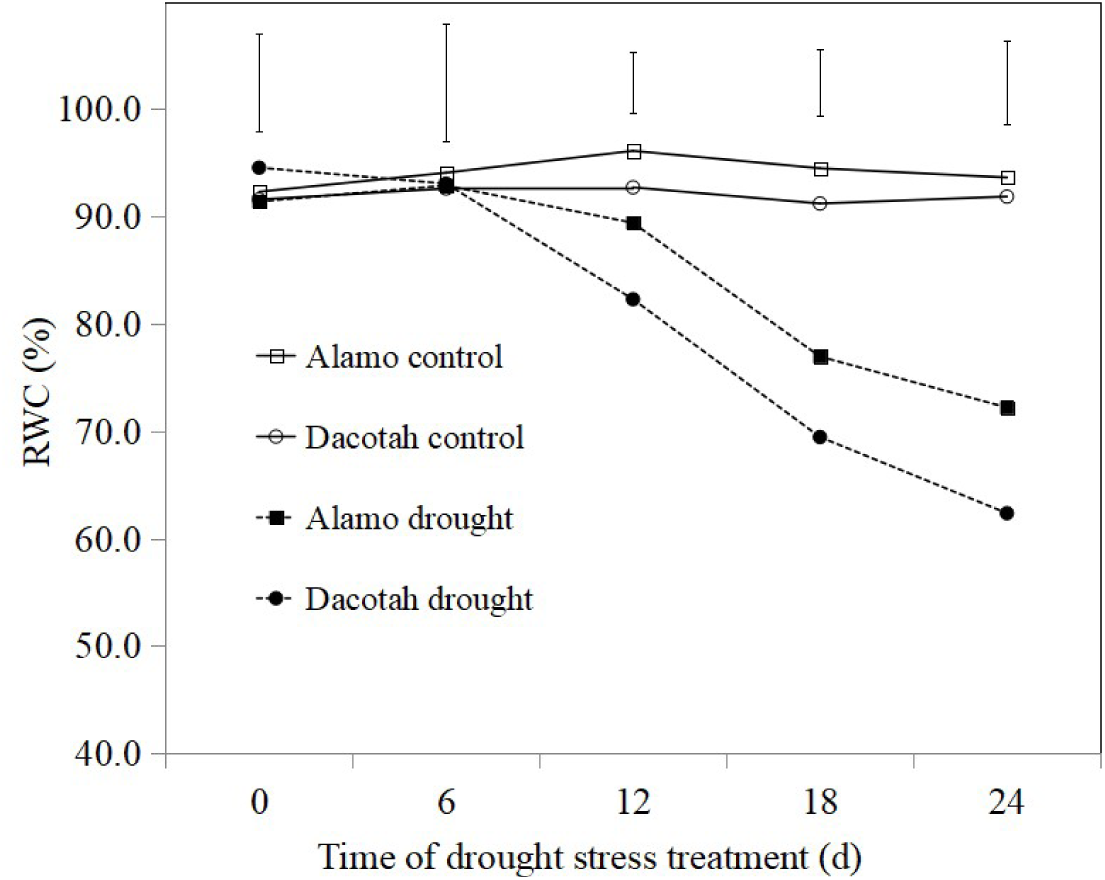
Effects of drought stress on relative water content (RWC) of two switchgrass cultivars Alamo and Dacotah. Vertical bars indicate LSD value (P=0.05).

**Fig. 4.**
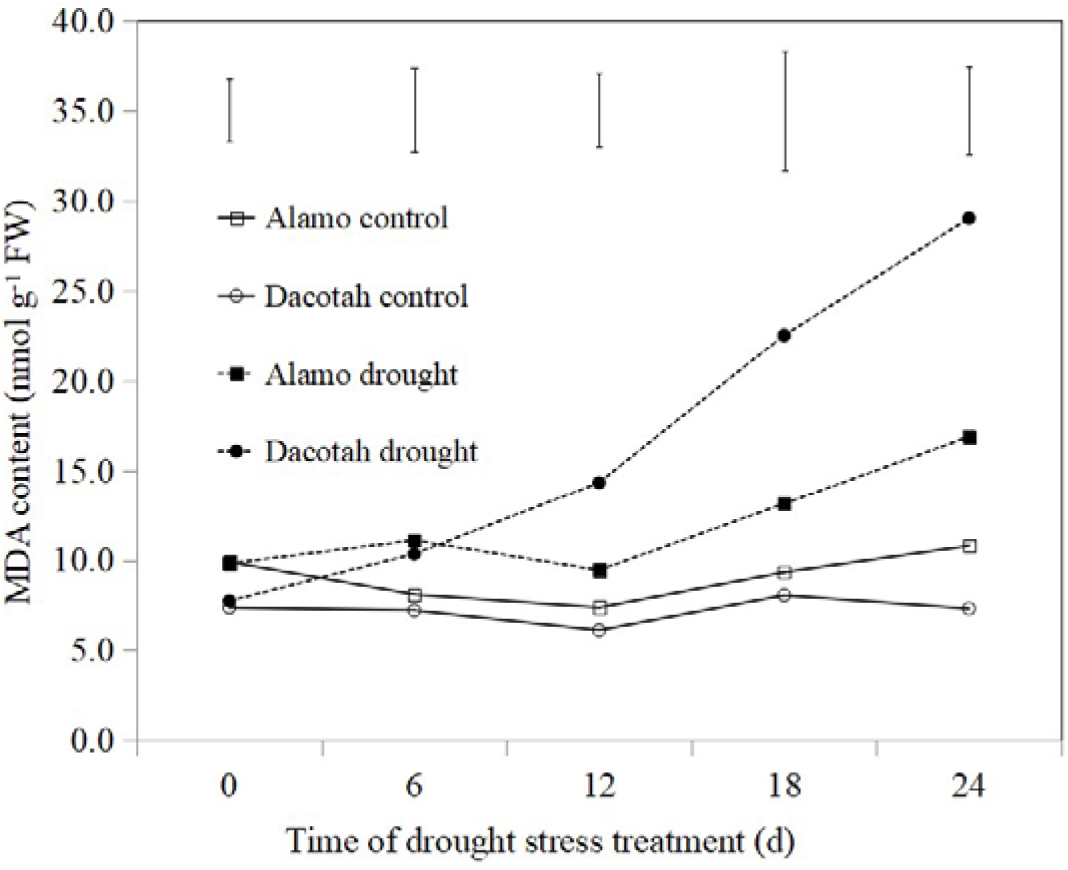
Effects of drought stress on malondialdehyde (MDA) of two switchgrass cultivars Alamo and Dacotah. Vertical bars indicate LSD value (P=0.05).

### Effects of drought stress on antioxidant enzymes activity

SOD activity of both cultivars increased with the increasing duration of drought stress, with a larger extent in drought tolerant Alamo than sensitive Dacotah (Fig. 5). A significant increase in SOD was observed at 18 d and 24 d of drought treatment for two cultivars compred to their control. At 24 d of drought treatment, SOD of Alamo increased to 150.0 μmol min^-1^ mg^-1^ protein, 3.65 times of control, SOD of Dacotah increased to 98.3 μmol min^-1^ mg^-1^ protein, 2.64 times of control, SOD of Alamo treatment was 1.53 times higher than Dacotah treatment.

**Fig. 5.**
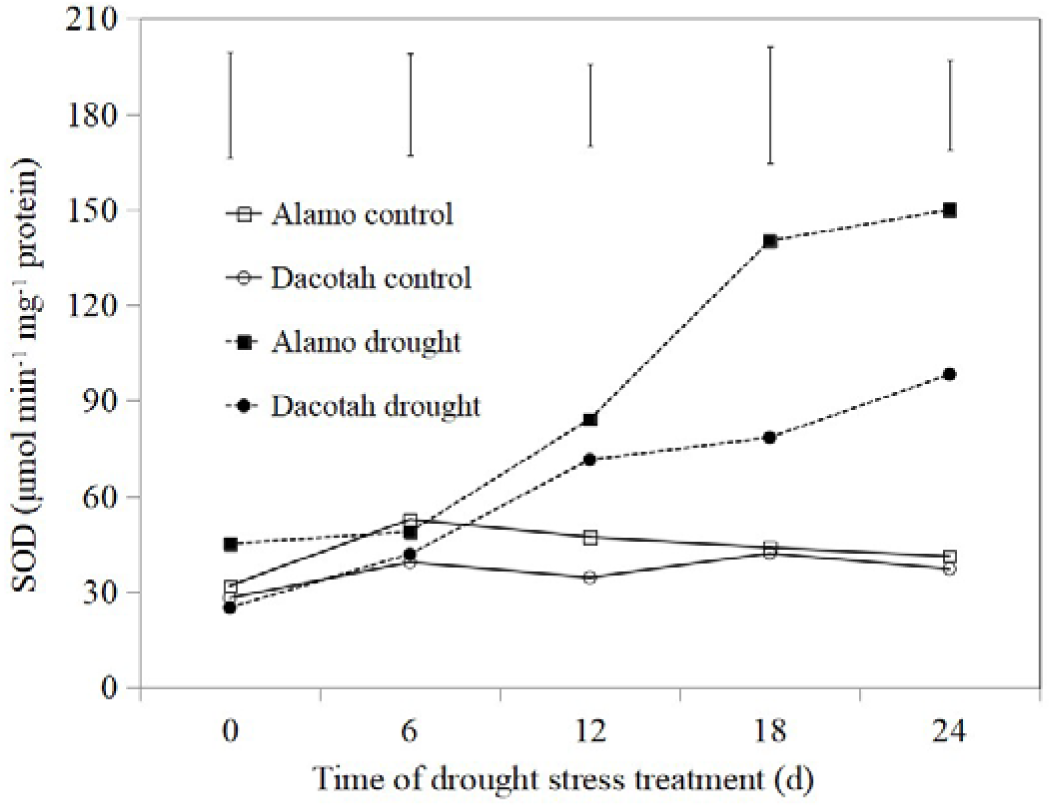
Effects of drought stress on superoxide dismutase (SOD) of two switchgrass cultivars Alamo and Dacotah. Vertical bars indicate LSD value (P=0.05).

CAT activity were progressively enhanced in the two cultivars with duration of drought stress compared to their control (Figs. 6), Alamo had greate CAT activity than Dacotah at 12, 18 and 24 d of drought treatment, CAT of Alamo treatment at 24 d was 1.29 times higher than Dacotah treatment at 24 d.

**Fig. 6.**
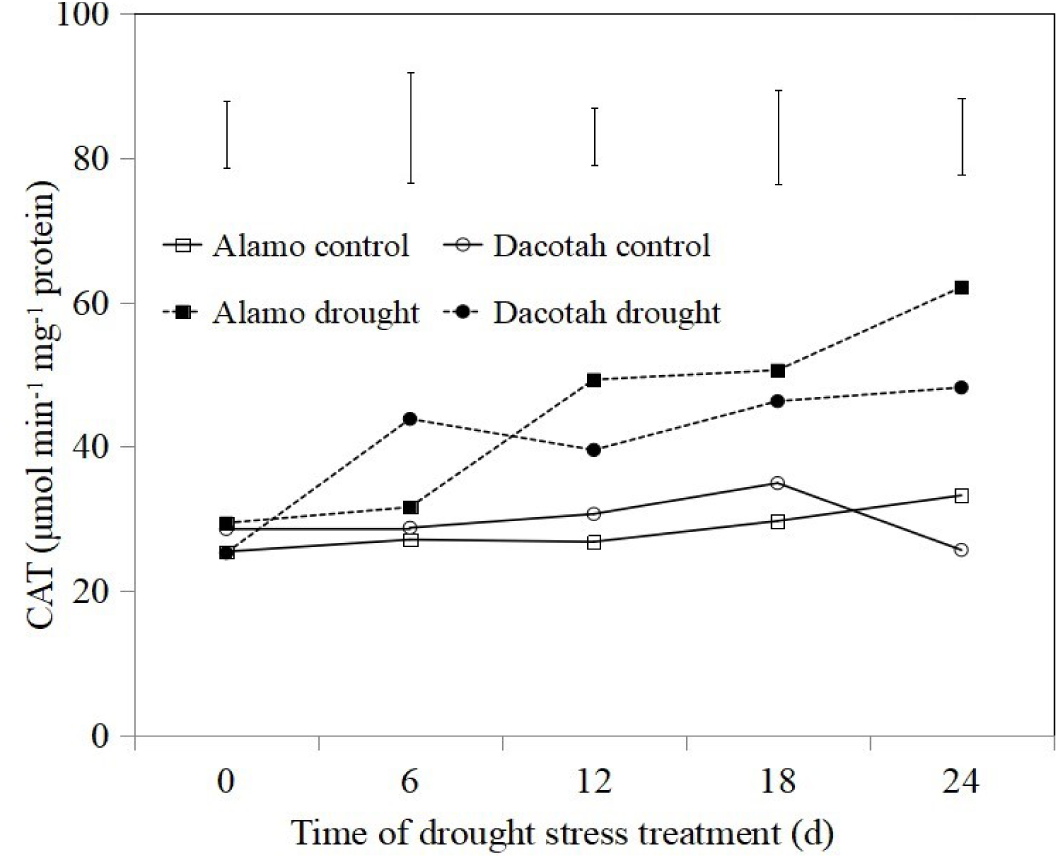
Effects of drought stress on ascorbate catalase (CAT) of two switchgrass cultivars Alamo and Dacotah. Vertical bars indicate LSD value (P=0.05).

APX had the same trend with SOD and CAT (Fig. 7). APX activity of both cultivars increased with the increasing duration of drought stress, the increase of Alamo was significant higher than Dacotah especially at 24 d of drought treatment. At 24 d of drought treatment, APX of Alamo increased to 73.4 μmol min^-1^ mg^-1^ protein, 2.01 times of control, APX of Dacotah increased to 49.8 μmol min^-1^ mg^-1^ protein, 1.54 times of control, SOD of Alamo treatment was 1.47 times higher than Dacotah treatment.

**Fig. 7.**
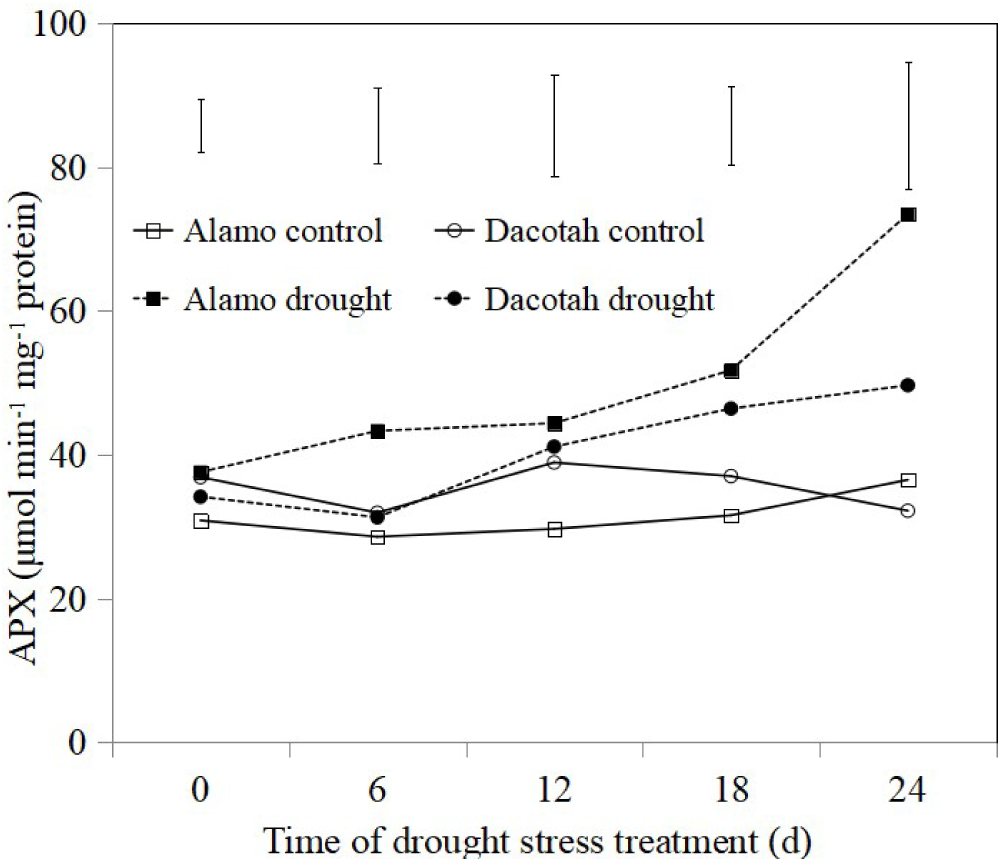
Effects of drought stress on ascorbate peroxidase (APX) of two switchgrass cultivars Alamo and Dacotah. Vertical bars indicate LSD value (P=0.05).

### Effects of drought stress on antioxidant isozyme

Our activity staining visualized four SOD isozymes (SOD1-SOD4) in two cultivars (Fig. 8). Alamo and Dacotah increased SOD1 abundance under drought stress compared to control, both cultivars increased SOD2-SOD3 abundance under drought stress compared to control, however, only Dacotah decreased SOD4 abundance under drought stress compared to control. Alamo had relative higher SOD3-SOD4 abundance in response to drought stress compared to Dacotah. Only one isoform of APX was identified in two cultivars (Fig. 9), both cultivars increased APX1 abundance under drought stress compared to control, Alamo had relative higher APX1 abundance in response to drought stress when compared to Dacotah.

**Fig. 8.**
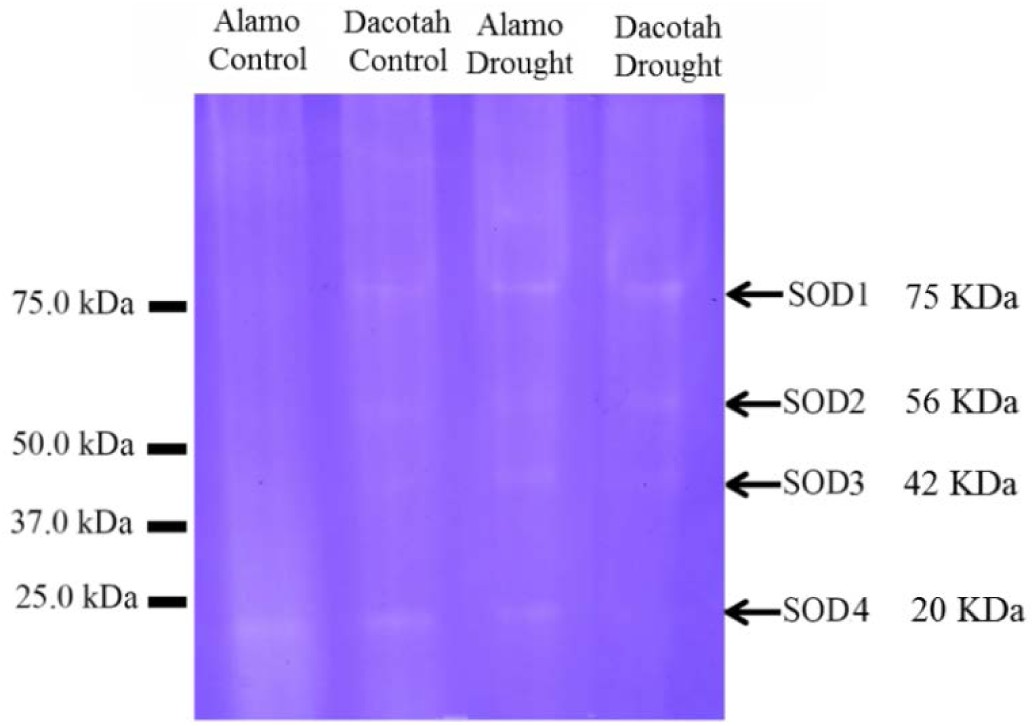
Changes in superoxide dismutase (SOD) isoforms of two switchgrass cultivars Alamo and Dacotah under control (well-watered) and drought stress conditions (24 d). Equal amounts (15 μL) were loaded in each lane.

**Fig. 9.**
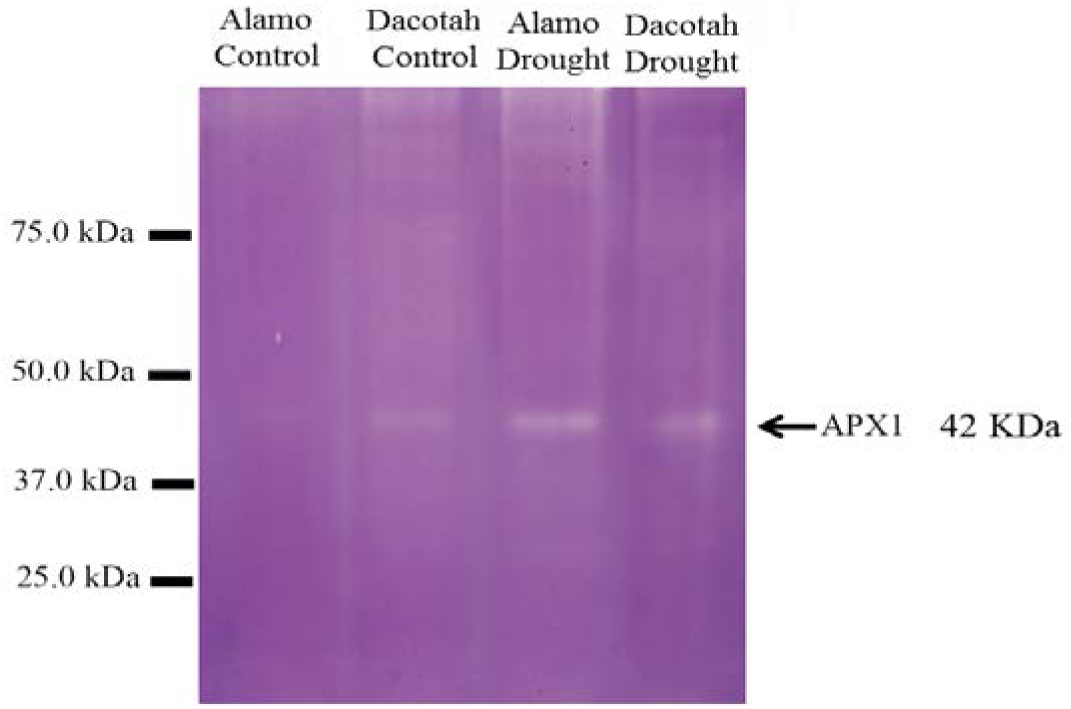
Changes in ascorbate peroxidase (APX) isoforms of two switchgrass cultivars Alamo and Dacotah under control (well-watered) and drought stress conditions (24 d). Equal amounts (15 μL) were loaded in each lane.

### Effects of drought stress on dehydrin

Western bolot showed that almost no dehydrin accumulation observed in control of both cultivars, however, drought stress significantly increased dehydrin accumulation in both cultivars at 24 d of drought treatment, Alamo had higher abundance of 53 KDa and 18 KDa dehydrin than Dacotah, no dehydrin accumulation of 18 KDa was found in Dacotah (Fig. 10).

**Fig. 10.**
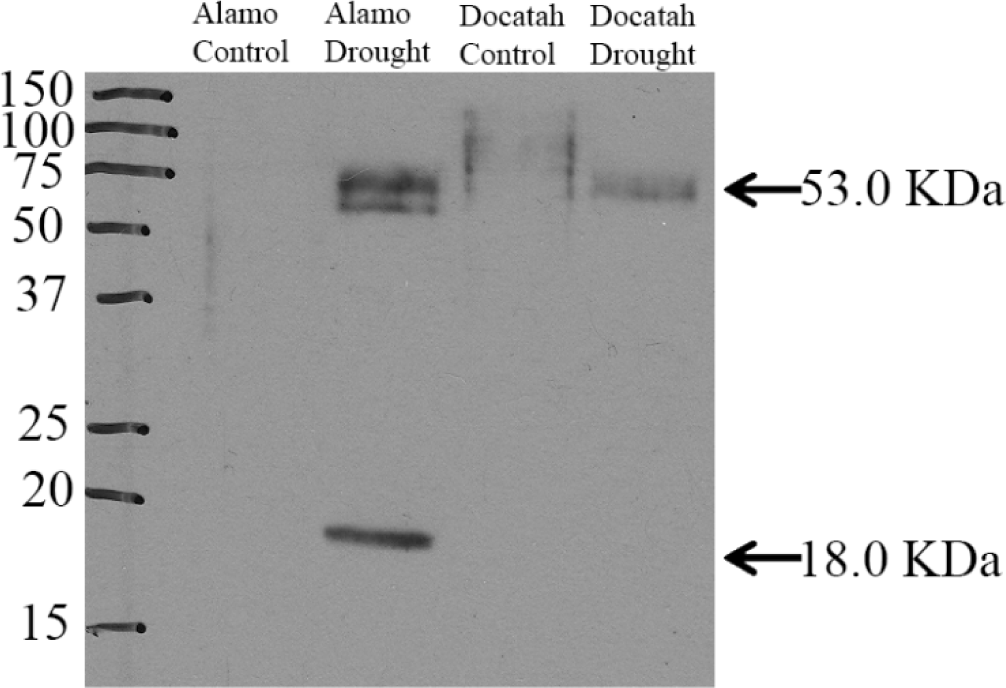
Immunoblots of dehydrin expression in two switchgrass cultivars Alamo and Dacotah under drought stress (24 d).

### qRT-PCR of antioxidant, dehydrin, heat shock protein (HSP) and aquaporin

qRT-PCR results showed that, the level of *PvSOD1*, *PvERD1*, *PvHSP90* and *PvPIP1;5* mRNA were significantly increased in both cultivars under drought treatment compared to their control, the level of *PvCAT1*, *PvAPX2* were significantly increased only in Alamo at 24 d of drought treatment compared to the control instead of Dacotah (Fig. 11). Whereas the increase of *PvSOD1* mRNA level in Dacotah was significant higher than Alamo at 24 d of drought stress, no significant difference in level of *PvHSP90* mRNA between Alamo and Dacotah at 24 d of drought stress.

**Fig. 11.**
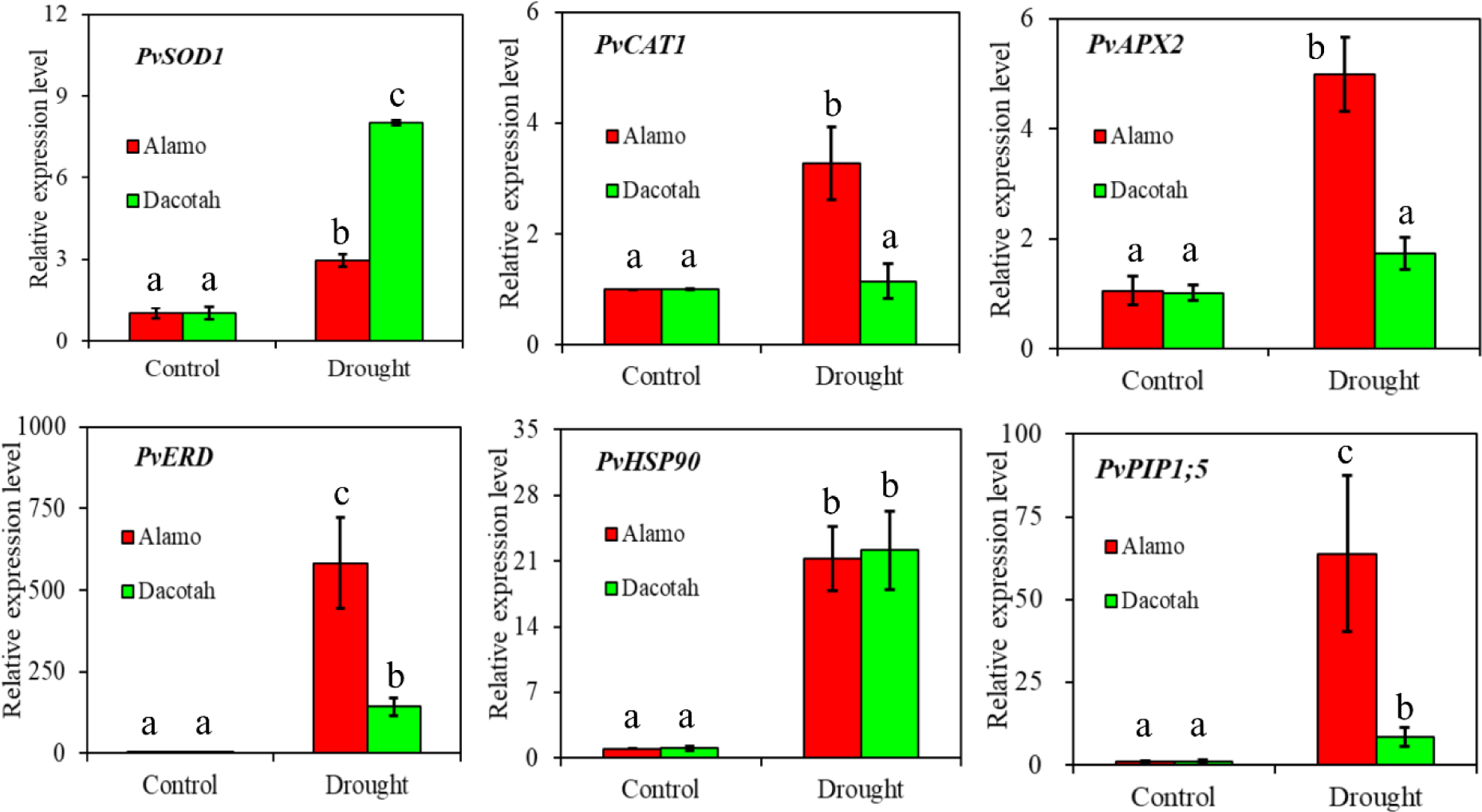
Relative transcript levels of *PvAPX2*, *PvCAT1*, *PvERD1*, *PvHSP90*, *PvSOD1* and PvPIP1;5 genes of two switchgrass cultivars (drought-tolerant Alamo and drought-sensitive Dacotah) under well-watered and drought stress (24 d). Each bar represents the mean of three independent replicates with standard error. Different letters of a-c indicates the statistic difference at 0.05 level.

## Discussion

Our previous study found that, Alamo ranked #4 in drought tolerance and Dacotah ranked #48 among 49 switchgrass lines, Alamo is a representative of good drought tolerant cultivars and Dacotah is a representative of drought sensitive ones [26]. The results of this study agreed with our previous study, showed that drought stress caused cellular and leaf damage to switchgrass as indicated by decreased leaf RWC, increased EL and MDA. Alamo had higher RWC, *PvPIP1;5* relative gene expression level, lower EL and MDA content than Dacotah at 24 d of drought treatment, indicating that drought stress resulted in more severe damage to cell membranes in Docatah relative to Alamo at 24 d of drought treatment.

When plants are subjected to water deficit, drought tolerant cultivars have better Tr and g_s_ and gas exchange and less oxidative stress, changes in antioxidants genes expression and activities may improve antioxidant defense system and ROS scavenging [36]. SOD is regarded as the key enzyme in the ROS scavenger system because it catalyzes superoxide free radical dismutation into H_2_O_2_ and O_2_, which is the first step of scavenging ROS [37]. APX and POD scavenge H_2_O_2_ and reduce ROS toxicity. In this study, we choose relative drought-tolerant cultivar Alamo, and relative drought-sensitive cultivar Dacotah to study the drought tolerance mechanisms of drought associated with antioxidant defense. Our results showed that, drought stress increase SOD, CAT and APX in two cultivars especially at 18 d and 24 d treatment. Alamo had significant higher SOD, CAT, APX activities and *PvAPX2*, *PvCAT1* relative gene expression level than Dacotah. R Khanna-Chopra *et al*. [38] found that drought-resistant wheat maintained favorable water relations and lower H_2_O_2_ accumulation during severe water stress conditions due to systematic increase of antioxidants such as SOD, APX, CAT [39]. Pallavi Sharma *et al*. [11] reported that drought stress caused oxidative damage in rice species as exhibited by an increase in O^•–^_2_ and H_2_O_2_, but drought tolerant rice maintained higher antioxidant enzyme activities (SOD, APX) compared with drought-sensitive rice. These results indicated that antioxidant defense plays an important role in improvement of drought tolerance in plants, drought tolerant cultivars may have greater ROS-scavenging capacity to suppress ROS-induced injury during abiotic stress.

Antioxidant isozymes are known to play a vital role in plant drought tolerance. In this study, antioxidant isozyme diversity of two switchgrass cultivars were investigated to identify band profiles as biochemical marker for the drought tolerance. Drought stress increased abundance of SOD1-SOD3 and APX1 isozyme in two cultivars at 24 d of drought stress, however, Dacotah had relatively lower SOD4 and APX1 abundance when compared to Alamo at 24 d of drought stress. The results of this study indicate that the drought tolerant cultivar Alamo had greater ability to scavenge ROS than the drought sensitive Dacotah. Our results agreed with the previous studies, which showed that various drought-tolerant plants present different isozyme patterns during drought stress conditions [40, 41].

Drought induced-accumulation of dehydrin proteins has been associated with drought tolerance in many plant species [42–44]. In this study, two dehydrin polypeptides (53 KDa and 18 KDa) were detected in Alamo under drought stress conditions at 24 d of drought treatment. Interestingly, only one dehydrin polypeptides (53 KDa) was detected in Dacotah, that could be caused by the different drought tolerant ability between the two cultivar under drought stress. Consistently, Alamo had significant higher *PvERD1* relative gene expression level than Dacotah.

In summary, drought stress caused damage to switchgrass as evidenced by decreased RWC, increased leaf EL and MDA content due to drought stress treatment. Alamo had relatively higher RWC, lower EL and less MDA content when compared to Dacotah at 24 d of drought stress. SOD, APX and CAT activities increased during drought stress at 18 d and 24 d. Alamo had higher SOD, APX and CAT activities, greater abundance of SOD1-SOD3 and APX1 isozyme than Dacotah under drought stress. Alamo had greater abundance of two dehydrin polypeptides (53 KDa and 18 KDa) under drought stress conditions at 24 d than Dacotah. qRT-PCR showed that antioxidant gene *PvAPX2*, *PvCAT1*, dehydrin gene *PvERD1*, and aquaporin gene *PvPIP1;5* instead of heat shock protein gene *PvHSP90* and *PvSOD1*, are the key genes contribute to the drought tolerance of Alamo, although Alamo had higher SOD activity and SOD isozyme abundance. Our results suggest antioxidant and dehydrin expression are associated with drought tolerance in switchgrass. Our results also suggest that selection and use of cultivars with greater antioxidant enzyme activities and more abundant isozymes under drought stress may be a practical approach to improve switchgrass drought tolerance.

## Acknowledgements

This project was supported by grants from the “Hainan Provincial Natural Science Foundation of China (No. 318MS101)”, “Central Public-interest Scientific Institution Basal Research Fund for Chinese Academy of Tropical Agricultural Sciences (No. 1630032017085/1630032018020)”, “China Agriculture Research System (No. CARS-22-Z11)”, “The Investigation of Forage Germplasms in South China (2017FY100600)” and “The U.S. Department of Agriculture (XZ and BZ) the Office of Science (BER), U.S. Department of Energy (KC, RB, XZ, and BZ)”.

## References

1. Kaur G, Asthir B. Molecular responses to drought stress in plants. Biol Plantarum. 2017;61(2):201–9.

2. Van Bavel CHM. Water Relations of Plants and Soils. Soil Science. 1996;161(4):257–60.

3. Krame P. Plant water relations. Academic Press, New York. 1983.

4. Sharma P, Jha AB, Dubey RS, Pessarakli M. Reactive Oxygen Species, Oxidative Damage, and Antioxidative Defense Mechanism in Plants under Stressful Conditions. Journal of Botany. 2012.

5. Bailey-Serres J, Mittler R. The roles of reactive oxygen species in plant cells. Plant Physiol. 2006;141(2):311.

6. Noctor G, Mhamdi A, Foyer CH. The roles of reactive oxygen metabolism in drought: not so cut and dried. Plant Physiol. 2014;164(4):1636–48.

7. Mittler R. ROS Are Good. Trends in Plant Science. 2017;22(1):11–9. doi: 10.1016/j.tplants.2016.08.002.

8. Mignolet-Spruyt L, Xu E, Idänheimo N, Hoeberichts FA, Mühlenbock P, Brosché M, et al. Spreading the news: subcellular and organellar reactive oxygen species production and signalling. J Exp Bot. 2016;67(13):3831–44. doi: 10.1093/jxb/erw080. PubMed PMID: 26976816.

9. Bray EA. Plant responses to water deficit. Trends in Plant Science. 1997;2(2):48–54. doi: 10.1016/S1360-1385(97)82562-9.

10. Prochazkova D, Sairam RK, Srivastava GC, Singh DV. Oxidative stress and antioxidant activity as the basis of senescence in maize leaves. Plant Sci. 2001;161(4):765–71.

11. Sharma PD, R.S. Drought Induces Oxidative Stress and Enhances the Activities of Antioxidant Enzymes in Growing Rice Seedlings. Plant Growth Regulation. 2005;46(3):209–21.

12. Uzilday B, Turkan I, Sekmen AH, Ozgur R, Karakaya HC. Comparison of ROS formation and antioxidant enzymes in Cleome gynandra (C4) and Cleome spinosa (C3) under drought stress. Plant Sci. 2012;182:59–70. Epub 2011/11/29.

13. Li Z, Peng Y, Huang B. Alteration of Transcripts of Stress-Protective Genes and Transcriptional Factors by γ-Aminobutyric Acid (GABA) Associated with Improved Heat and Drought Tolerance in Creeping Bentgrass (*Agrostis stolonifera*). Int J Mol Sci [Internet]. 2018; 19(6).

14. Ren J, Sun LN, Zhang QY, Song XS. Drought Tolerance Is Correlated with the Activity of Antioxidant Enzymes in Cerasus humilis Seedlings. Biomed Res Int. 2016;2016:9851095. doi: 10.1155/2016/9851095. PubMed PMID: 27047966.

15. Sen A, Alikamanoglu S. Characterization of drought-tolerant sugar beet mutants induced with gamma radiation using biochemical analysis and isozyme variations. J Sci Food Agric. 2014;94(2):367–72. doi: 10.1002/jsfa.6393. PubMed PMID: 24037781.

16. Sanyal RP, Samant A, Prashar V, Misra HS, Saini A. Biochemical and functional characterization of OsCSD3, a novel CuZn superoxide dismutase from rice. Biochemical Journal. 2018. doi: 10.1042/bcj20180516.

17. Graether SP, Boddington KF. Disorder and function: a review of the dehydrin protein family. Frontiers in plant science [Internet]. 2014 2014; 5:[576 p.].

18. Hu L, Wang Z, Du H, Huang B. Differential accumulation of dehydrins in response to water stress for hybrid and common bermudagrass genotypes differing in drought tolerance. J Plant Physiol. 2010;167(2):103–9.

19. Lv A, Fan N, Xie J, Yuan S, An Y, Zhou P. Expression of CdDHN4, a Novel YSK2-Type Dehydrin Gene from Bermudagrass, Responses to Drought Stress through the ABA-Dependent Signal Pathway. Frontiers in Plant Science. 2017;8(748). doi: 10.3389/fpls.2017.00748.

20. Lopez CG, Banowetz GM, Peterson CJ, Kronstad WE. Dehydrin Expression and Drought Tolerance in Seven Wheat Cultivars. Crop Science. 2003;43:577–82. doi: 10.2135/cropsci2003.5770.

21. Han B, Kermode AR. Dehydrin-like proteins in castor bean seeds and seedlings are differentially produced in response to ABA and water-deficit-related stresses. J Exp Bot. 1996;47(7):933–9. doi: 10.1093/jxb/47.7.933.

22. McLaughlin SB, Adams Kszos L. Development of switchgrass (Panicum virgatum) as a bioenergy feedstock in the United States. Biomass and Bioenergy. 2005;28(6):515–35.

23. Jiang Y, Yao Y, Wang Y. Physiological response, cell wall components, and gene expression of switchgrass under short-term drought stress and recoveries. Crop Sci. 2012;52.

24. Hopkins AA, Taliaferro CM, Murphy CD, Christian DA. Chromosome Number and Nuclear DNA Content of Several Switchgrass Populations. Crop Science. 1996;36(5):1192–5.

25. Alexopoulou E, Sharma N, Papatheohari Y, Christou M, Piscioneri I, Panoutsou C, et al. Biomass yields for upland and lowland switchgrass varieties grown in the Mediterranean region. Biomass and Bioenergy. 2008;32(10):926–33.

26. Liu Y, Zhang X, Tran H, Shan L, Kim J, Childs K, et al. Assessment of drought tolerance of 49 switchgrass (*Panicum virgatum*) genotypes using physiological and morphological parameters. Biotechnology for Biofuels. 2015;8(1):152. doi: 10.1186/s13068-015-0342-8.

27. Hardin CF, Fu C, Hisano H, Xiao X, Shen H, Stewart CN, Jr., et al. Standardization of switchgrass sample collection for cell wall and biomass trait analysis. BioEnergy Research. 2013;6(2):755–62. doi: 10.1007/s12155-012-9292-1.

28. Marcum KB, Anderson SJ, Engelke MC. Salt gland ion secretion: a salinity tolerance mechanism among five zoysiagrass species. Crop Sci. 1998;38:806–10.

29. Barrs H, Weatherley P. A Re-Examination of the Relative Turgidity Technique for Estimating Water Deficits in Leaves. Australian Journal of Biological Sciences. 1962;15(3):413–28.

30. Giannopolitis CN RS. Superoxide dismutases, 1: Occurrence in higher plants [Corn, oats, peas]. Plant Physiol. 1977;59:309–14.

31. Chance B, Maehly AC. Assay of catalases and peroxidases. Methods in Enzymology. 2: Academic Press; 1955. p. 764–75.

32. Nakano Y AK. Hydrogen peroxide scavenged by ascorbate-specific peroxidase in spinach chloroplasts. Plant Cell Physiol. 1981;22(5): 867–80.

33. Dhindsa RS, Plumb-Dhindsa P, Thorpe TA. Leaf Senescence: Correlated with Increased Levels of Membrane Permeability and Lipid Peroxidation, and Decreased Levels of Superoxide Dismutase and Catalase. J Exp Bot. 1981;32(126):93–101.

34. Beauchamp C, Fridovich I. Superoxide dismutase: Improved assays and an assay applicable to acrylamide gels. Analytical Biochemistry. 1971;44(1):276–87.

35. Lopez-Huertas E, Corpas FJ, Sandalio LM, Del Rio LA. Characterization of membrane polypeptides from pea leaf peroxisomes involved in superoxide radical generation. The Biochemical journal. 1999;337 (Pt 3):531–6. Epub 1999/01/23. PubMed PMID: 9895298; PubMed Central PMCID: PMCPMC1220006.

36. Bartosz G. Oxidative stress in plants. Acta Physiol Plant. 1997;19(1):47–64. doi: 10.1007/s11738-997-0022-9.

37. Bowler C, Montagu MV, Inze D. Superoxide Dismutase and Stress Tolerance. Annual Review of Plant Physiology and Plant Molecular Biology. 1992;43(1):83–116. doi: doi:10.1146/annurev.pp.43.060192.000503.

38. R Khanna-Chopra DS. Acclimation to drought stress generates oxidative stress tolerance in drought-resistant than -susceptible wheat cultivar under field conditions. Environmental & Experimental Botany. 2007;60(2):276–83.

39. Selote DS, Khanna-Chopra R. Antioxidant response of wheat roots to drought acclimation. Protoplasma. 2010;245(1-4):153–63. Epub 2010/06/19. doi: 10.1007/s00709-010-0169-x. PubMed PMID: 20559854.

40. Zhang X, Ervin EH, Liu Y, Hu G, Shang C, Fukao T, et al. Differential Responses of Antioxidants, Abscisic Acid, and Auxin to Deficit Irrigation in Two Perennial Ryegrass Cultivars Contrasting in Drought Tolerance. Journal of the American Society for Horticultural Science. 2015;140(6):562–72.

41. Luo N, Yu X, Nie G, Liu J, Jiang Y. Specific peroxidases differentiate Brachypodium distachyon accessions and are associated with drought tolerance traits. Ann Bot-London. 2016;118(2):259–70. doi: 10.1093/aob/mcw104.

42. Hassan NM, El-Bastawisy ZM, El-Sayed AK, Ebeed HT, Nemat Alla MM. Roles of dehydrin genes in wheat tolerance to drought stress. J Adv Res. 2015;6(2):179–88. doi: 10.1016/j.jare.2013.11.004. PubMed PMID: 25750752.

43. Verma G, Dhar YV, Srivastava D, Kidwai M, Chauhan PS, Bag SK, et al. Genome-wide analysis of rice dehydrin gene family: Its evolutionary conservedness and expression pattern in response to PEG induced dehydration stress. Plos One [Internet]. 2017 2017; 12(5):[e0176399 p.].

44. Kumar M, Lee S-C, Kim J-Y, Kim S-J, Aye SS, Kim S-R. Over-expression of dehydrin gene, OsDhn1, improves drought and salt stress tolerance through scavenging of reactive oxygen species in rice (*Oryza sativa* L.). J Plant Biol. 2014;57(6):383–93. doi: 10.1007/s12374-014-0487-1.

